# Distinct roles of α- and β-tubulin C-terminal tails for ciliary function as revealed by a CRISPR/Cas9 mediated gene editing in *Chlamydomonas*

**DOI:** 10.1101/2023.02.14.528553

**Authors:** Tomohiro Kubo, Yuma Tani, Haru-Aki Yanagisawa, Masahide Kikkawa, Toshiyuki Oda

## Abstract

α- and β-tubulin have an unstructured glutamate-rich region at their C-terminal tails (CTT). The function of this region in cilia/flagella is still unclear, except that glutamates in CTT act as the sites for posttranslational modifications that affect ciliary motility. A unicellular alga *Chlamydomonas* possesses only two α-tubulin genes and two β-tubulin genes, each pair encoding an identical protein. This simple gene organization may enable a complete replacement of the wild-type tubulin with its mutated version. Here, using CRISPR/Cas9, we generated mutants expressing tubulins with modified CTTs. We found that the mutant whose four glutamate residues in the α-tubulin CTT have been replaced by alanine almost completely lacked polyglutamylated tubulin and displayed paralyzed cilia. In contrast, the mutant lacking the glutamate-rich region of the β-tubulin CTT assembled short cilia without the central apparatus. This phenotype is similar to the mutants harboring a mutation in a subunit of katanin, whose function has been shown to depend on the β-tubulin CTT. Therefore, our study reveals distinct and important roles of α- and β-tubulin CTT in the formation and function of cilia.

**Summary statement:** *Chlamydomonas* mutants were produced by CRISPR/Cas9 mediated gene editing to investigate ciliary function of tubulin C-terminal tails (CTTs). We found that α- and β-tubulin CTTs are essential for ciliary motility and assembly.

## Introduction

A tubulin molecule consists of an ∼400 amino-acid core and an unstructured ∼20 amino-acid C-terminal tail (CTT) abundant in glutamate residues (Nogales et al., 1998). In microtubules, the C-terminal region protrudes from protofilaments and affects various aspects of microtubule function: tubulin polymerization (Serrano et al., 1984), microtubule severing (Roll-Mecak and Vale, 2008), kinetochore dynamics (Miller et al., 2008), and interactions with motor proteins (Wang and Sheetz et al., 2000; Sirajuddin et al., 2014) and microtubule-associated proteins (Hinrichs et al., 2012). However, the function of tubulin CTT in cilia and flagella (interchangeable terms), whose abnormality causes various human diseases collectively called the “ciliopathy,” is still obscure.

Several kinds of posttranslational modifications (PTMs), including polyglutamylation and polyglycylation, occur in the tubulin CTT. Polyglutamylation, a modification on the γ-carboxyl group of glutamate residues, forms side chains of up to 20 glutamate units (Eddé et al., 1990; Rüdiger et al., 1992; Audebert et al., 1994; Redeker et al., 2005). We and others have previously shown that this modification regulates ciliary motility (Kubo et al., 2010; Suryavanshi et al., 2010; Ikegami et al., 2010) by affecting an axonemal structure called the nexin-dynein regulatory complex (Kubo et al., 2012; 2017). Another kind of tubulin modification, polyglycylation, competes with polyglutamylation for the same glutamate residues and generates side chains of up to 34 glycyl units (Redeker et al., 1994; 2005). Polyglycylation was shown to be involved in the stability and maintenance of ependymal cilia (Bosch Grau et al., 2013) and primary cilia (Rocha et al., 2014). Recently, this modification was also found to affect the motility of mouse sperm by modulating the power generation of axonemal dyneins (Gadadhar et al., 2021). In mammals, seven out of the 13 members of the tubulin tyrosine ligase-like (TTLL) protein family possess glutamylation activity (Janke et al., 2005; Dijk et al., 2007), and three other members possess glycylation activity (Wloga et al., 2009; Ikegami et al., 2009). In cilia, the levels of glutamylation and glycylation affect each other; namely, the lack of glutamylation causes the increase in glycylation and vice versa (Redeker et al., 2005). However, physiological significance of this anticorrelation between the two PTMs for ciliary functions is still unclear.

We used *Chlamydomonas reinhardtii*, which possesses only two α-tubulin genes (*TUA1* and *TUA2*) and two β-tubulin genes (*TUB1* and *TUB2*), each pair encoding an identical protein (Silflow et al., 1985; Figure 1A, 5A). Such a small number of tubulin genes and species variation is a feature not seen in mammalian cells, which normally have multiple genes for both α- and β-tubulins. The simple gene organization of tubulins in *Chlamydomonas* may enable complete replacements of the wild-type tubulin with mutated versions by gene editing. In this study, using a CRISPR/Cas9, we generated strains harboring mutations in the CTT of either α- or β-tubulin. For α-tubulin, we generated mutants in which the four glutamate residues in the CTT were replaced by alanine. For β-tubulin, we produced mutants lacking variable lengths of the CTT region where multiple glutamate residues are clustered. The α-tubulin mutant named *TUA1(4A)* without four glutamate residues (E445, E447, E449, E450) was found to completely lack polyglutamylation and ciliary motility. On the other hand, the β-tubulin CTT mutant *TUB2(Δ6E)* without six glutamates (E435, E437, E439, E440, E441, E442) was found to lack glycylation and assemble only short cilia without the central-pair microtubules. Our study thus reveals distinct and important roles of α- and β-tubulin CTTs for ciliary function.

**Figure 1.**
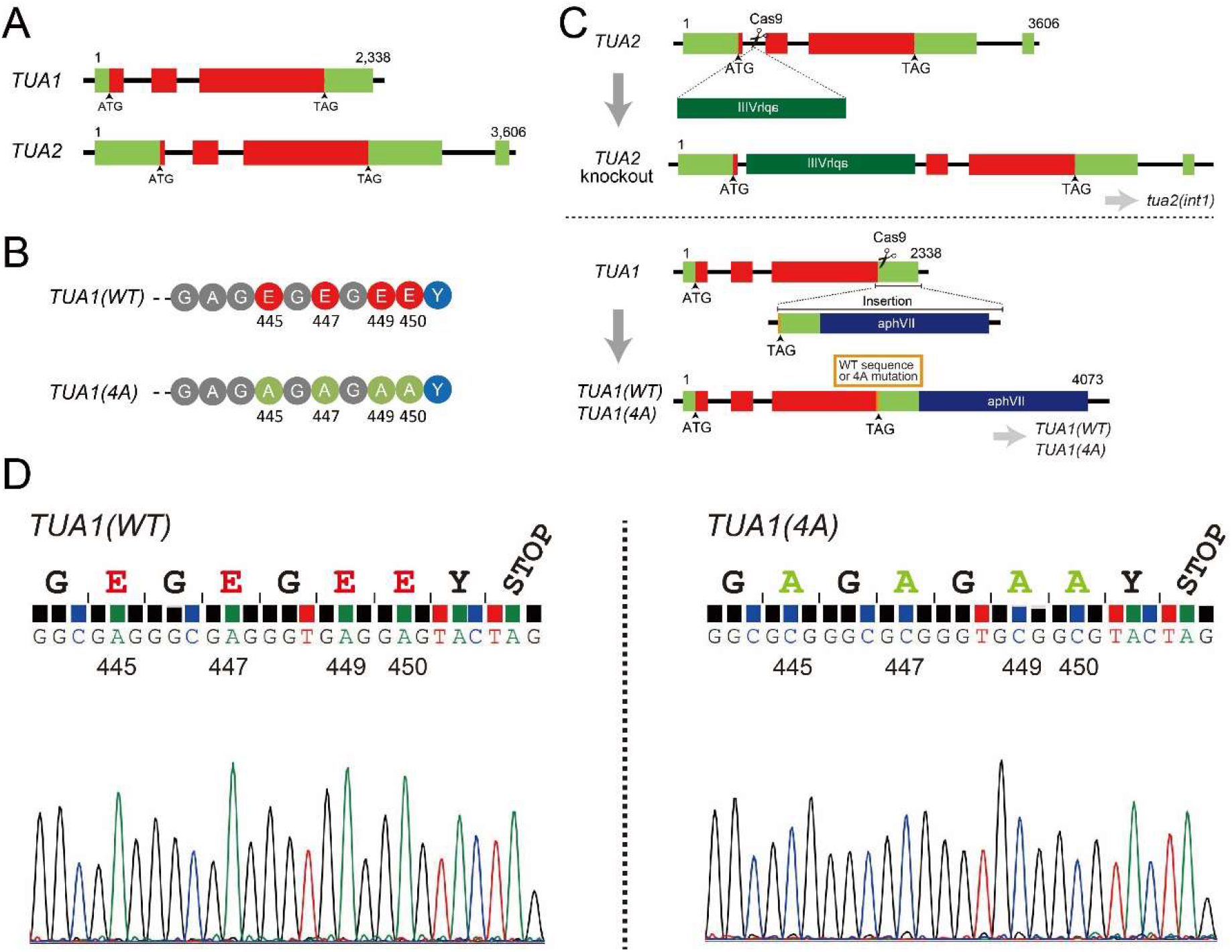
Generation of a mutant possessing mutations in the α-tubulin C-terminal tail. (A) The structures of *TUA1* (Cre03.g190950) and *TUA2* (Cre04.g216850) genes. Both genes encode identical proteins. (B) Candidate sites of α-tubulin polyglutamylation. These four glutamates (E445, E447, E449, and E450; shown in red) were replaced with aspartates (shown in green). (C) Production of *TUA1(WT)* and *TUA1(4A)* by CRISPR/Cas9 mediated gene editing. First, a paromomycin-resistant gene cassette was inserted into *TUA2* gene of wild type to generate *tua2(int1)* (upper panel). The mutant *tua2(int1)* was found to express only *TUA1* for α-tubulin. Secondly, *TUA1* C-terminus region of *tua2(int1)* was replaced with a sequence encoding wild-type or mutated *TUA1* C-terminus, 3’UTR, and a hygromycin resistant-gene to produce *TUA1(WT)* and *TUA1(4A)* (lower panel). (D) Genomic sequences encoding the C-terminal tail of TUA1 in the *TUA1(WT)* and *TUA1(4A)* strains.

## Results

### Generation of a mutant lacking glutamate residues in the α-tubulin CTT

We set out to produce an α-tubulin CTT mutant lacking the four glutamate residues (E445, E447, E449, and E450; Figure 1B) predicted as glutamylation and glycylation sites (Redeker et al., 2005). Because either one of the two α-tubulin genes of *Chlamydomonas, TUA1* (Cre03.g190950) and *TUA2* (Cre04.g216850) (Figure 1A), can complement the lack of the other (Fromherz et al., 2004; Kato-Minoura et al., 2020), we used a strain lacking the *TUA2* gene and replaced all the four glutamates with alanine in the TUA1 protein.

First, a *TUA2-*knockout strain was generated. A paromomycin-resistant gene cassette was inserted into the intron 1 of *TUA2* in the wild-type strain by a CRISPR/Cas9 mediated gene editing (Figure 1C, upper panel). Insertion was detected by PCR and verified by sequencing (Supplemental Figure 1). Semi-quantitative RT-PCR showed that *TUA2* was not expressed whereas *TUA1* expression was nearly doubled in a transformant named *tua2(int1)* (not shown) as in other α-tubulin mutants generated by insertional-mutagenesis (Kato-Minoura et al., 2020). This indicates that α-tubulin is expressed exclusively from *TUA1* in the *tua2(int1)* cell. Next, we introduced a specific donor DNA into the *TUA1-*coding sequence in *tua2(int1)*. The donor DNA encodes a C-terminal sequence of the TUA1 protein with or without the mutations (Figure 1B), the 3’ UTR of *TUA1*, and a hygromycin-resistant gene cassette (Figure 1C, lower panel; Supplemental Figure 2). PCR screening identified several candidates for both the control and the *TUA1* mutant, which were subsequently selected by sequencing of *TUA1* gene for the control and a mutant with four “GAG” to “GCG” mutations reflecting glutamate to alanine replacements (Figure 1D). Representative strains were chosen and named “*TUA1(WT)*” for the control and “*TUA1(4A)*” for the *TUA1* mutant with the glutamate replacements.

**Figure 2.**
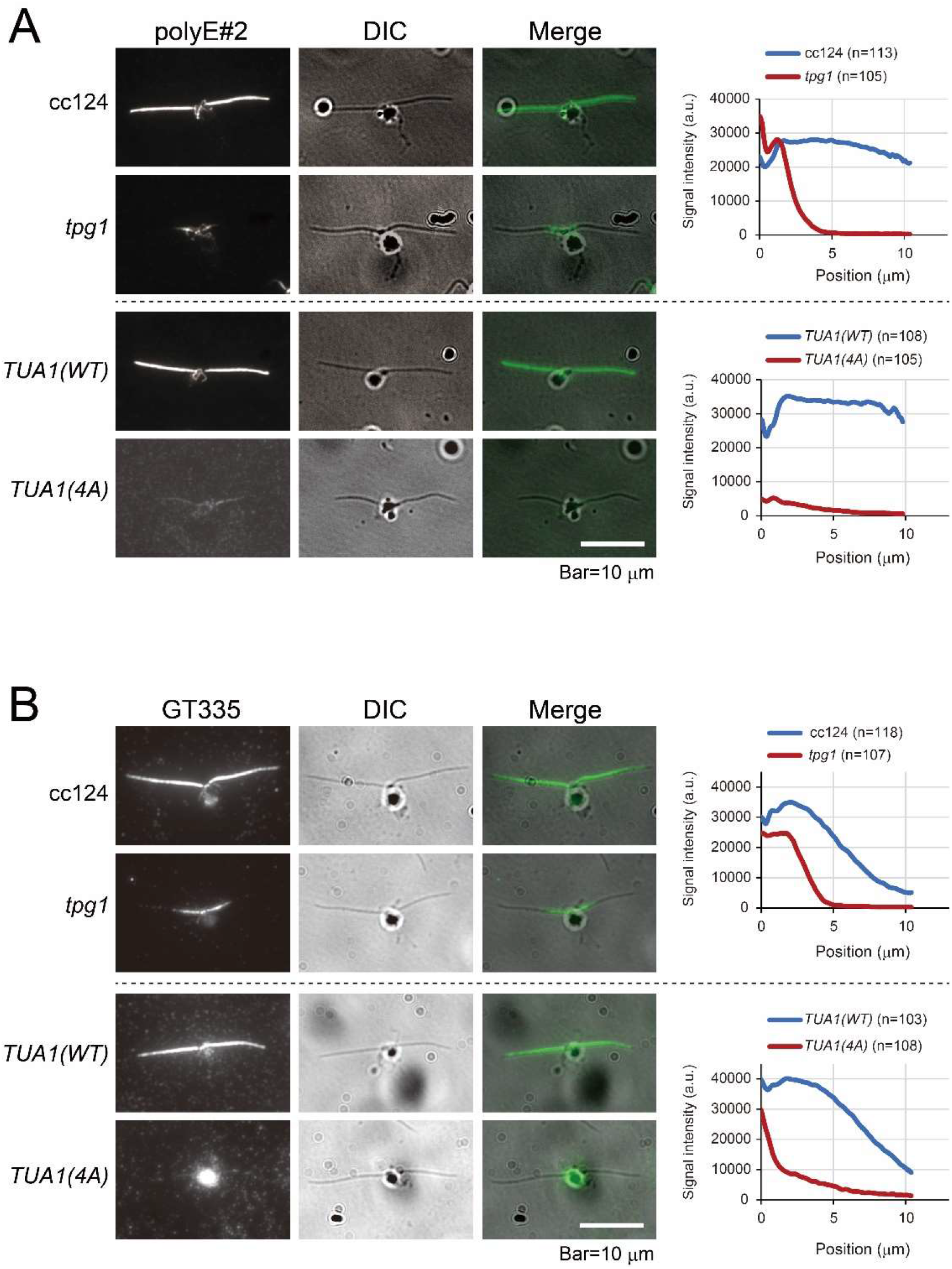
The mutant *TUA1(4A)* completely lacks α-tubulin glutamylation. Immunostaining of nuclear-flagellar apparatus (NFAp) of wild type (cc124), *tpg1, TUA1(WT)*, and *TUA1(4A)* using (A) polyE#2 and (B) GT335 antibodies. One cilium each on at least 100 cells was investigated and the average signal intensity was obtained.

### Tubulin polyglutamylation occurs mostly on the α-tubulin CTT in *Chlamydomonas*

We performed indirect immunofluorescence microscopy of nuclear-flagellar apparatus (NFAp) isolated from wild type (cc124), *tpg1, TUA1(WT)*, and *TUA1(4A)* to examine the level of polyglutamylation. The mutant *tpg1* lacks TTLL9 polyglutamylase and thereby has a reduced level of axonemal tubulin polyglutamylation (Kubo et al., 2010). PolyE#2, an antibody that recognizes side chains of three or more glutamates (Kubo and Oda, 2017), showed uniform staining of both wild-type and *TUA1(WT)* axonemes along the entire length (Figure 2A). The *tpg1* axoneme had a signal only in the proximal region, consistent with our previous study (Figure 2A; Kubo et al., 2010). In contrast to wild-type and *tpg1* NFAps, the *TUA1(4A)* NFAps displayed very faint fluorescence (Figure 2A). GT335, an antibody that recognizes glutamate side chains of any length including monoglutamylation, showed a staining pattern different from the polyE#2 pattern (Figure 2B). The wild-type and *TUA1(WT)* axonemes showed a signal along the length with decreasing intensity towards the tip (Figure 2B). The *tpg1* axoneme displayed a signal only in one third of the proximal region. In contrast, the *TUA1(4A)* axoneme showed essentially no signal (Figure 2B). However, for an unknown reason, the nuclei had intense staining in *TUA1(4A)*.

Western blotting of the axonemes supported the immunostaining observation (Figure 3B). In contrast to the wild-type axonemes, which were intensely polyglutamylated, the *tpg1* and *TUA1(4A)* axonemes were much less glutamylated. Absence of both polyE and GT335 signals in the *TUA1(4A)* axonemes suggests that polyglutamylation occurs mostly on the α-tubulin CTT. This is consistent with our previous conclusion that β-tubulin is poorly glutamylated in the *Chlamydomonas* axoneme (Kubo et al., 2010).

**Figure 3.**
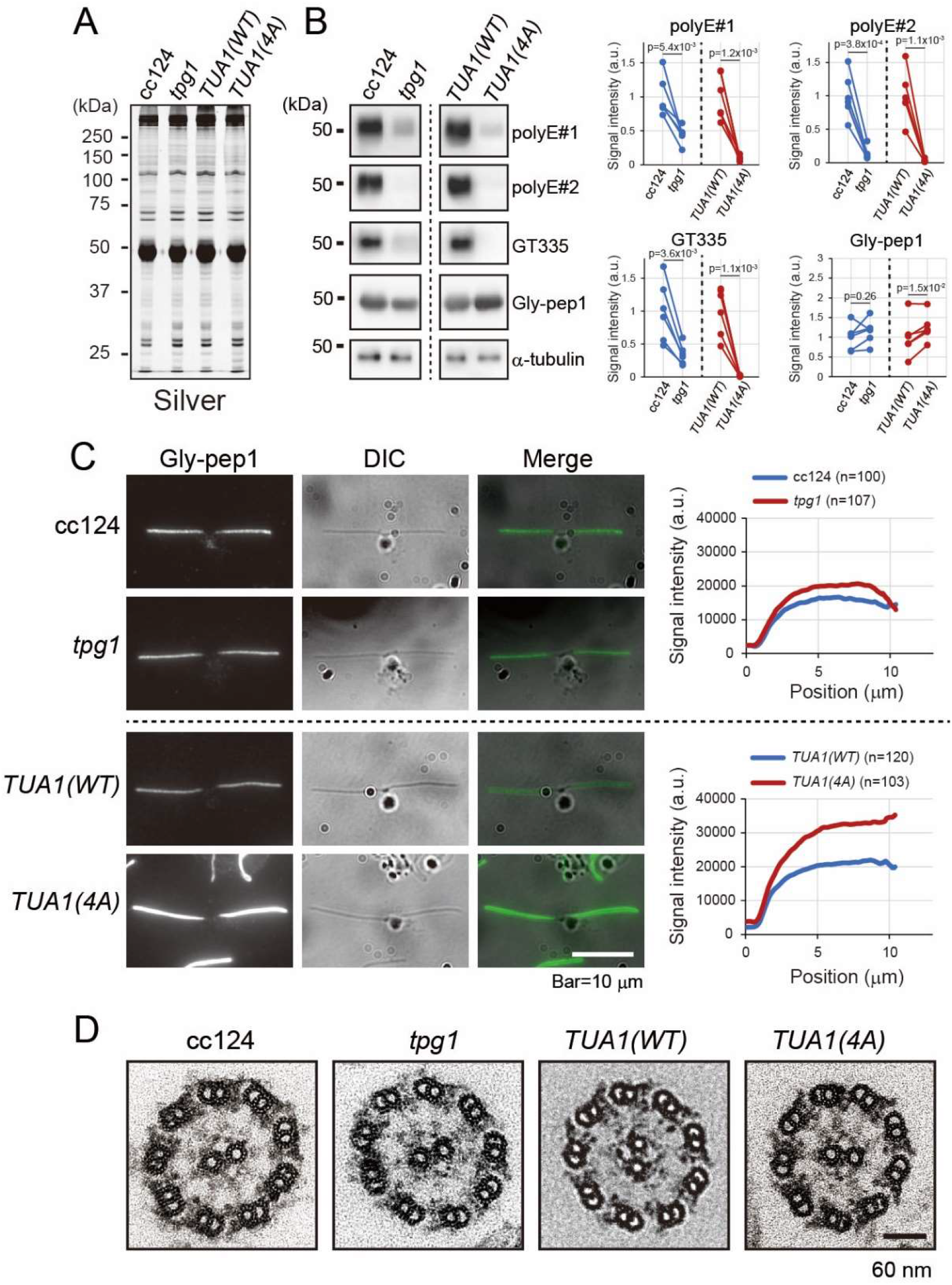
The *TUA1(4A)* axoneme has normal structure with increased glycylation. (A) Silver-stained gel and (B) Western blotting of cc124, *tpg1, TUA1(WT)*, and *TUA1(4A)* axonemes. For the Western blotting, six different blots of each pair were statistically analyzed by t-test. (C) Indirect immunofluorescence microscopy using Gly-pep1 antibody. One cilium each on at least 100 different cells was investigated and average signal intensity was obtained. (D) Cross sections of the axoneme observed by transmission electron microscope.

Indirect immunofluorescence microscopy with an antibody (Gly-pep1) recognizing mono- and bi-glycylated tubulin demonstrated that glycylated tubulin was present along the full length of both wild-type and *TUA1(WT)* axonemes but was absent in their basal bodies and cytoplasmic microtubules (Figure 3C). Interestingly, the level of glycylation increased by ∼1.5 times in the *TUA1(4A)* axoneme compared to that of *TUA1(WT)* (Figure 3B, 3C). This result is in accordance with a previous report that mutation in the α-tubulin CTT causes an increase in β-tubulin glycylation, implicating a “cross-talk” between α- and β-tubulin polymodification (Redeker et al., 2005). It must be noted, however, that an antibody recognizing tubulin with long polyglycylated side chains failed to detect signals by Western blotting in any of the axonemal samples used (not shown). As in human cilia (Bré et al., 1996; Rogowski et al., 2009), glycylated side chains most likely do not significantly elongate in *Chlamydomonas* cilia. Indeed, *Chlamydomonas* genome does not have a gene encoding a protein homologous to TTLL10, an enzyme that elongates glycyl side chains (Kubo and Oda, 2019).

### α-tubulin CTT affects ciliary motility possibly through modulation of the inter-doublet friction in the axoneme

Although the *TUA1(4A)* axoneme had an apparently normal protein composition (Figure 3A) and structure (Figure 3D), the cells of this strain were severely defective in motility and showed only some irregular movements (Figure 4A). In contrast, the *tpg1* cells can swim at a reduced velocity (Figure 4A). This suggests that residual polyglutamylation or the glutamate residues of the α-tubulin CTT in the *tpg1* axoneme is important for ciliary motility.

**Figure 4.**
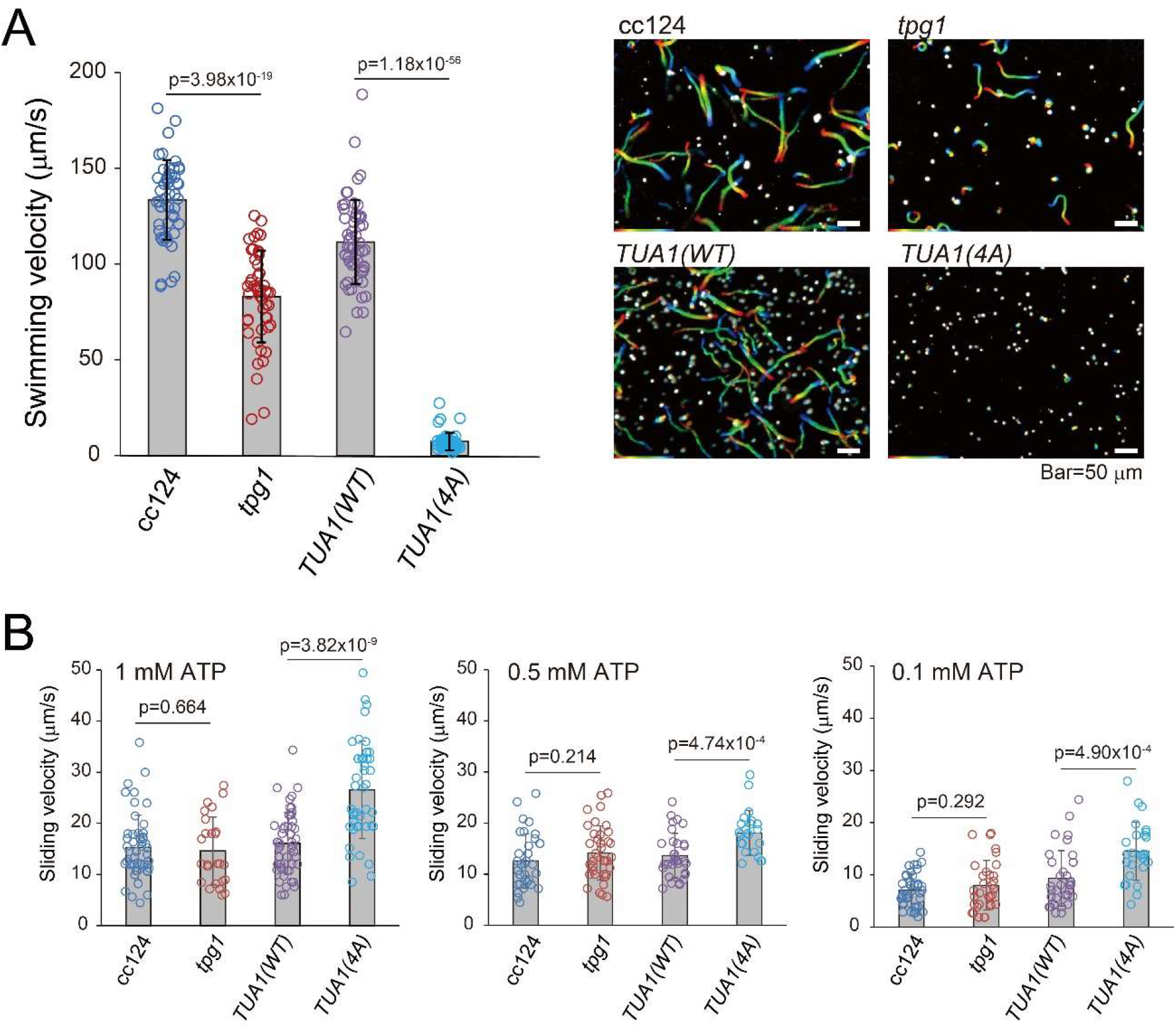
The mutant *TUA1(4A)* almost completely lacks ciliary motility but shows extremely fast axonemal microtubule sliding. (A) Swimming velocities (left) and trajectories (right; 1 s exposure) of wild type (cc124), *tpg1, tua2(int1), TUA1(WT)*, and *TUA1(4A)*. (B) Axonemal sliding velocities of cc124, *tpg1, TUA(WT)*, and *TUA1(4A)* in the presence of 1, 0.5, and 0.1 mM ATP.

To further explore the motility of *TUA1(4A)* cilia, we performed sliding disintegration assay, wherein fragmented axonemes are treated with a protease in the presence of ATP to induce microtubule sliding (Okagaki and Kamiya, 1986; Kurimoto and Kamiya, 1991). Strikingly, nonmotile *TUA1(4A)* axoneme did display sliding disintegration and the velocity of microtubule sliding was much higher than the velocity in the axonemes of wild type, *TUA1(WT)*, or *tpg1* (Figure 4B). The *tpg1* mutation has also been shown to produce faster microtubule sliding in the axoneme lacking outer-arm dyneins (Kubo et al., 2010; 2012). Therefore, the result of *TUA1(4A)* supports the idea that the negative charges of the tubulin C-terminus region increase the inter-microtubule friction (or some inter-doublet binding force) occurring in the axoneme (Kubo et al., 2010; 2012). Together with the fact that *TUA1(4A)* is unable to swim, it is most likely that the polyglutamylation or glutamates in the α-tubulin CTT are necessary for proper force generation of dynein required for the regular ciliary beating.

### Generation of mutants lacking C-terminal tail of β-tubulin

We next investigated the function of the β-tubulin CTT by truncating its glutamate-rich region. *Chlamydomonas* has two β-tubulin genes, *TUB1* (Cre12.g542250) and *TUB2* (Cre12.g549550) (Figure 5A), encoding an identical polypeptide with nine glutamate residues in their CTT (Figure 5B). We first generated a *TUB1-*knockout mutant by inserting a paromomycin-resistant gene cassette into the *TUB1* gene of the wild-type strain (Figure 5C; upper panel). Genotyping and semi-quantitative RT-PCR revealed that *TUB1* expression is undetectable in the novel *tub1(int2)* mutant indicating that it expresses β-tubulin only from *TUB2* (not shown). We then introduced a donor DNA in the *TUB2* coding sequence of *tub1(int2)* to completely or partially truncate the glutamate-rich region of the β-tubulin CTT (Figure 5C; lower panel). We generated mutants lacking four or six glutamates by inserting a stop codon in the *TUB2* coding sequence near the 3’ UTR (Figure 5B, 5C). PCR and sequencing identified an insertion of the donor DNA encoding a stop codon, the 3’UTR of *TUB2*, and the hygromycin-resistant gene for the three transformants designated *TUB2(WT), TUB2(Δ4E)*, and *TUB2(Δ6E)*, respectively (Figure 5D). Despite repeated trials, we could not isolate a mutant lacking nine glutamates of β-tubulin CTT in the background of *tub1(int2)*, indicating that a mutant without nine glutamates in the β-tubulin CTT is lethal. All of the isolated strains proliferated normally in the TAP medium. The viability of *TUB2(Δ6E)* was unexpected, because *Tetrahymena* mutants whose β-tubulin glutamates were substituted in three out of the five positions in CTT (E437, E438, E439, E440, and E442) show a hypomorphic or lethal phenotype (Xia et al., 2000; Thazhath et al., 2002; Redeker et al., 2005).

**Figure 5.**
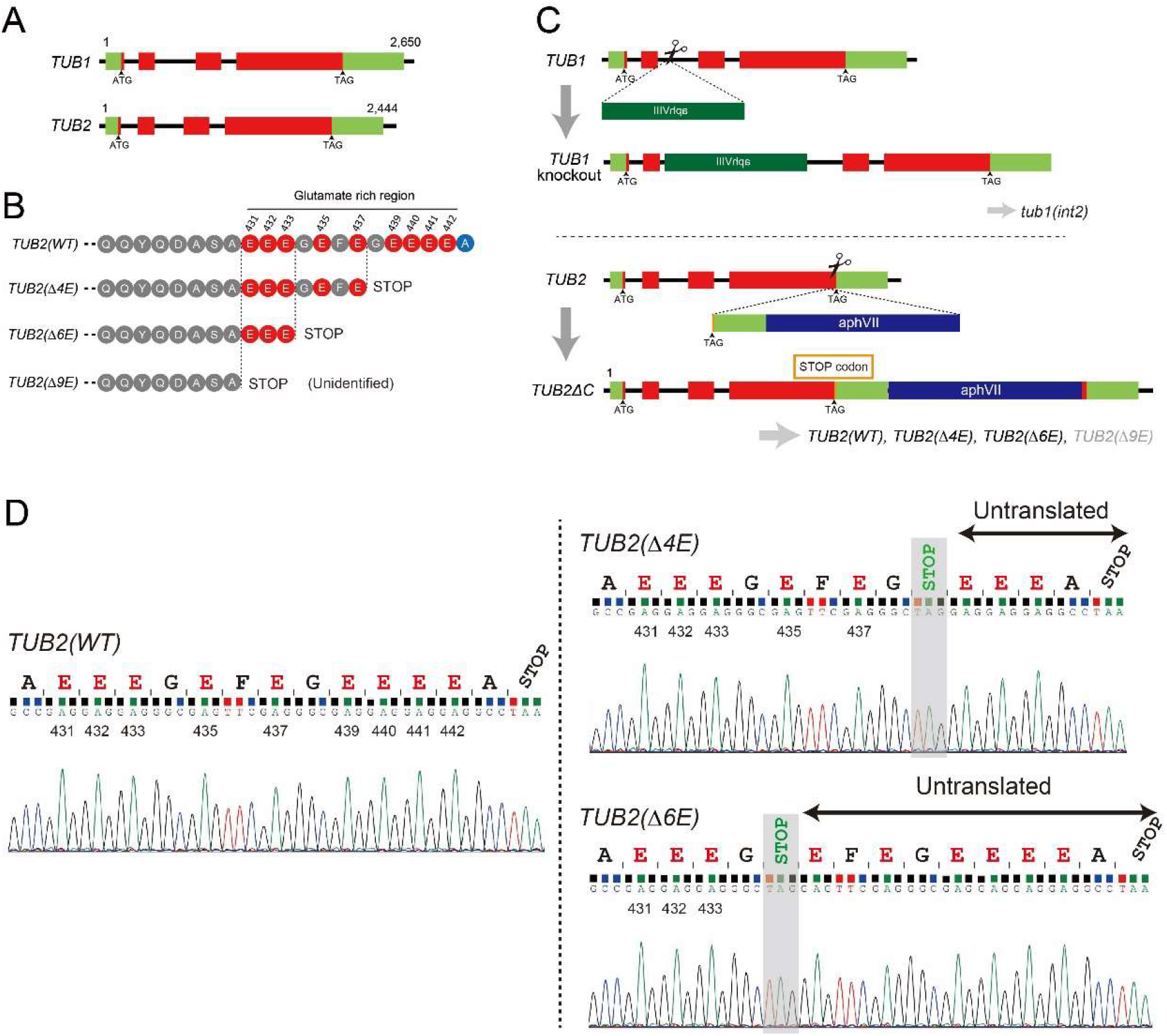
Generation of mutants lacking the β-tubulin C-terminal tail. (A) Structures of *TUB1* (Cre12.g542250) and *TUB2* (Cre12.g549550) genes. Both genes encode identical proteins. (B) Amino-acid sequence of the β-tubulin C-terminal tail. The glutamate region was partially abolished by introducing a stop codon. The mutant lacking nine glutamate residues was not identified. (C) Production of *tub1(int2)* and *TUB2(WT), TUB2(Δ4E), TUB2(Δ6E)*, and *TUB2(Δ9E)* by a CRISPR/Cas9 mediated gene editing. First, a paromomycin-resistant gene cassette was inserted into the *TUB1* gene of wild type to produce *tub1(int2)* mutant (upper panel). *tub1(int2)* expresses only *TUB2* for β-tubulin (not shown). Secondly, a stop codon and a hygromycin-resistant gene was inserted into *TUB2* of *tub1(int2)* (lower panel). Despite multiple trials, *TUB2(Δ9E)* was not identified. (D) Genomic sequences encoding the mutated TUB2 proteins in the *TUB2(WT), TUB2(Δ4E)*, and *TUB2(Δ6E)* strains.

### Mutants lacking the C-terminal tail of β-tubulin have short nonmotile cilia

Both *TUB2(Δ4E)* and *TUB2(Δ6E)* cells were severely impaired in motility (Figure 6A), suggesting that the β-tubulin CTT is also involved in ciliary function like the α-tubulin CTT. Most severely impaired was *TUB2(Δ6E)*, which did not display swimming at all (Figure 6A). This motility defect may well be due to the abnormal ciliary structure because the *TUB2(Δ6E)* cilia were about half the length of wild-type cilia (Figure 6B). In addition, *TUB2(Δ6E)* failed to regenerate cilia after pH-shock-induced ciliary detachment, while *TUB2(Δ4E)* showed almost normal ciliary regeneration (Figure 6C).

**Figure 6.**
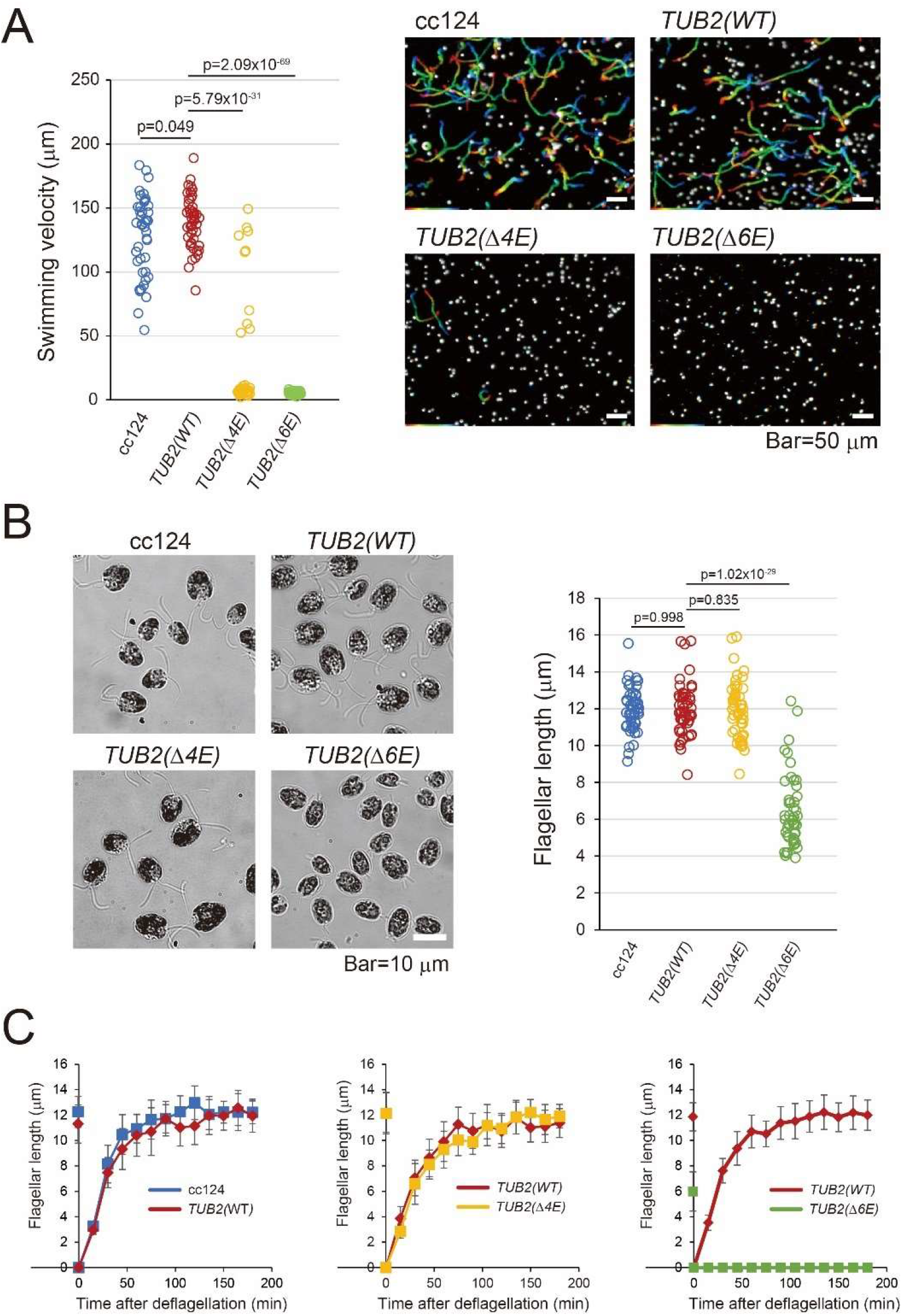
The mutant *TUB2(Δ6E)* displays short cilia without motility. (A) Swimming velocities (left) and trajectories (right; 1 s exposure) of wild type (cc124), *TUB2(WT), TUB2(Δ4E)*, and *TUB2(Δ6E)*. Ciliary lengths (B) and ciliary regeneration speeds (C) of wild type (cc124), *TUB2(WT), TUB2(Δ4E)*, and *TUB2(Δ6E)*.

### Tubulin glycylation takes place mostly on β-tubulin in *Chlamydomonas*

Western blotting using polyE and GT335 antibodies showed that the level of polyglutamylation is increased or unchanged in *TUB2(Δ4E)* and *TUB2(Δ6E)* axonemes (Figure 7A). In contrast, the glycylation levels of axonemes in these mutants were significantly decreased; in particular, the signal was almost completely missing in *TUB2(Δ6E)* (Figure 7A). Therefore, it is likely that glycylation mainly occurs on the β-tubulin CTT in *Chlamydomonas*.

**Figure 7.**
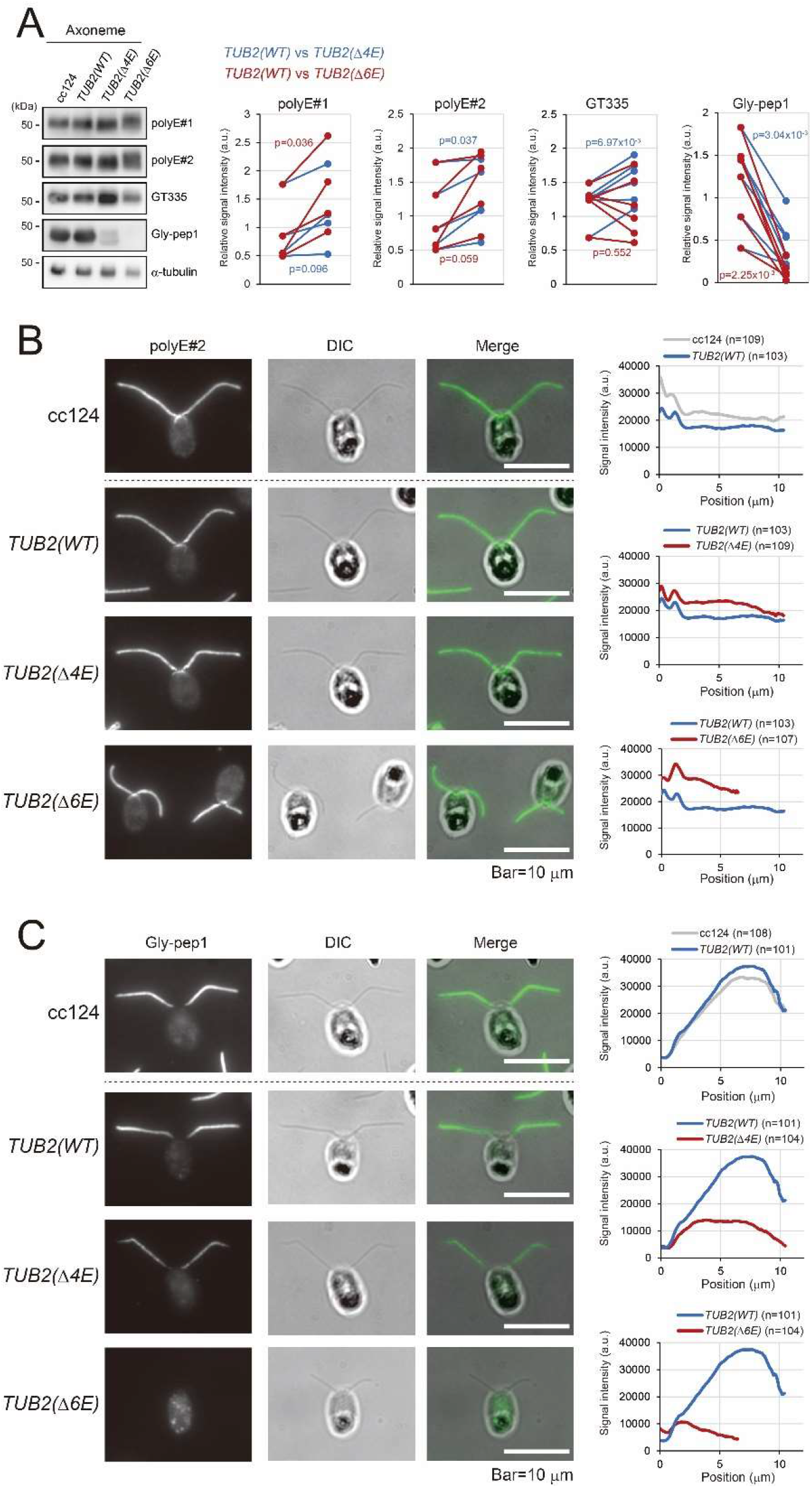
The mutants deficient in β-tubulin CTT lack tubulin glycylation. (A) Western blotting of the axonemes isolated from cc124, *TUB2(WT), TUB2(Δ4E)*, and *TUB2(Δ6E)* using the indicated antibodies. Signal intensities of the *TUB2(Δ4E)* and *TUB2(Δ6E)* axonemes were each compared to that of the *TUB2(WT)*. Immunostaining of cc124, *TUB2(WT), TUB2(Δ4E)*, and *TUB2(Δ6E)* using polyE#2 (B) or Gly-pep1 antibodies (C).

To determine the localization of axonemal glutamylation and glycylation, we performed immunostaining of NFAps. As expected, both *TUB2(Δ4E)* and *TUB2(Δ6E)* axonemes showed slightly increased polyE#2 staining (Figure 7B). Also as expected, while the NFAp of wild type and *TUB2(WT)* showed intense axonemal staining with the antibody recognizing glycylated tubulin (Gly-pep1), the NFAps of *TUB2(Δ4E)* and *TUB2(Δ6E)* had significantly decreased levels of the staining (Figure 7C).

### The *TUB2(Δ6E)* axoneme lacks the central apparatus

To explore the reason for the paralyzed cilia of *TUB2(Δ6E)*, we examined the cross-section images of the isolated axonemes by electron microscopy. The wild type (cc124), *TUB2(WT)*, and *TUB2(Δ4E)* axonemes had a normal “9+2” configuration (Figure 8A; Table 1). Interestingly, however, almost all of the *TUB2(Δ6E)* axoneme images lacked the two central-apparatus microtubules and, instead, contained electron-dense materials in the central lumen (Figure 8A). Several *Chlamydomonas* mutants are also known to lack the central apparatus. It is interesting to note that two central apparatus lacking mutants, *pf15* and *pf19*, have mutations in the subunits of katanin (Dymek et al., 2004; 2012), a protein complex that severs microtubules (McNally and Vale, 1993) depending on its interaction with the β-tubulin CTT (Zehr et al., 2020).

**Table 1.** The central-pair microtubules deficiency in the β-tubulin mutants

|  | cc124 |  | *TUB2(WT)* |  | *TUB2(Δ4E)* |  | *TUB2(Δ6E)* |  |
| --- | --- | --- | --- | --- | --- | --- | --- | --- |
|  | n | % | n | % | n | % | n | % |
| Central pair | 147 | 93.0 | 155 | 95.1 | 89 | 80.2 | 0 | 0.0 |
| Single microtubule | 10 | 6.3 | 4 | 2.5 | 19 | 17.1 | 1 | 1.0 |
| No Central pair | 1 | 0.6 | 4 | 2.5 | 3 | 2.7 | 57 | 54.3 |
| Electron dense material | 0 | 0.0 | 0 | 0.0 | 0 | 0.0 | 47 | 44.8 |
| Total | 158 | 100 | 163 | 100 | 111 | 100 | 105 | 100 |

**Figure 8.**
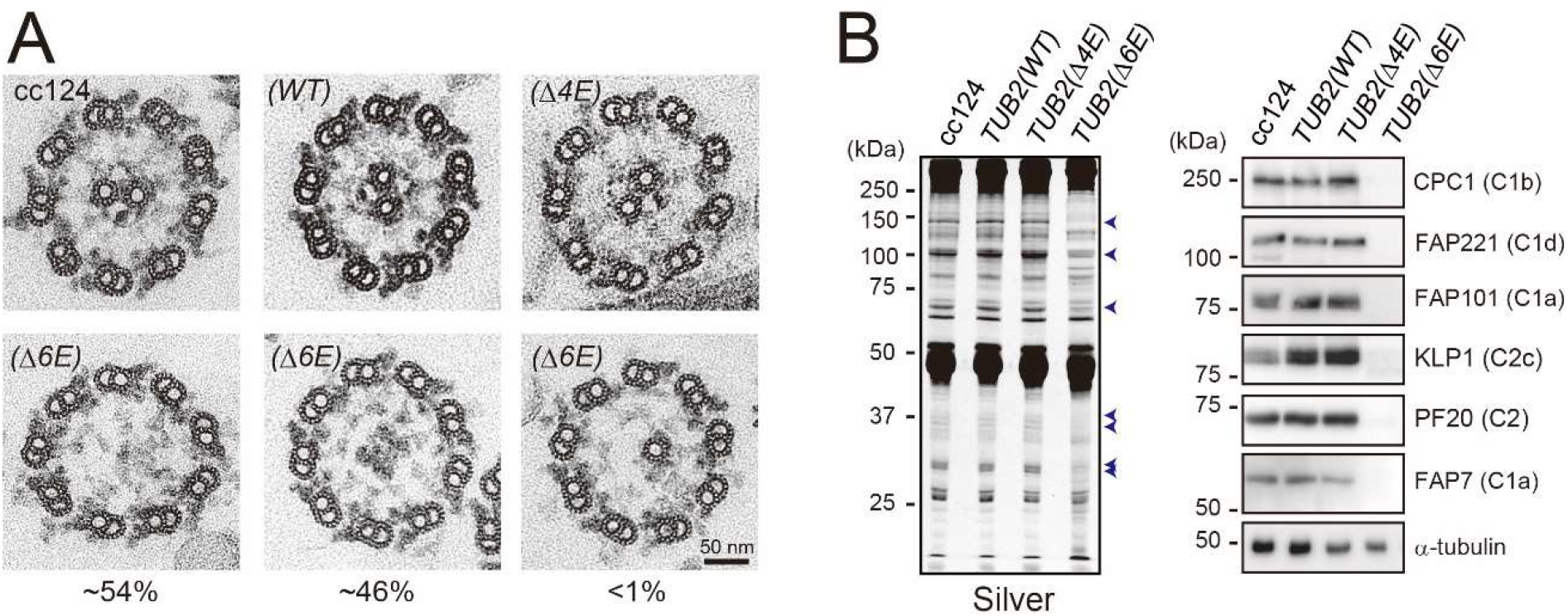
The *TUB2(Δ6E)* cilium lacks the central apparatus. (A) Cross section of the axoneme isolated from cc124, *TUB2(WT), TUB2(Δ4E)*, and *TUB2(Δ6E)* observed by transmission electron microscope. The mutant *TUB2(Δ6E)* lacks the central-pair microtubules. (B) (Left) Silver-stained gel of the isolated cilia. Bands decreased in the *TUB2(Δ6E)* were indicated by the arrowheads. (Right) Western blot of the isolated cilia using the antibodies against several central apparatus (CA) proteins. Predicted locations of the CA proteins in the C1 and C2 microtubules are based on Zhao et al. (2019) and Han et al. (2021).

The presence of central-apparatus proteins was examined in the axonemes from wild type, *TUB2(WT), TUB2(Δ4E)*, and *TUB2(Δ6E)* by a silver stained SDS-PAGE gel. While the *TUB2(Δ4E)* cilia showed a normal band pattern, the *TUB2(Δ6E)* cilia lacked several bands reflecting the deficiency of the central apparatus (Figure 8B, left). To assess the presence of the proteins that constitute the central-apparatus projections, polyclonal antibodies were raised against CPC1 (C1b projection; 205.1 kDa), FAP221 (C1d projection; 104 kDa), FAP101 (C1a projection; 85.7 kDa), KLP1 (C2c projection; 83 kDa), PF20 (C2 projection; 53 kDa), and FAP7 (C1a projection; 54.7 kDa) (Supplemental Figure 3) and used for Western blotting of ciliary samples. The results showed that the *TUB2(Δ6E)* cilia lack all the proteins examined (Figure 8B, right).

## Discussion

### Glutamate residues in both α- and β-tubulin CTTs are pivotal for ciliary motility

Our study suggests that glutamylation specifically occurs on α-tubulin whereas glycylation specifically occurs on β-tubulin in *Chlamydomonas* cilia. More specifically, some or all of the four glutamate residues (E445, E447, E449, and E450) in the α-tubulin CTT are likely the main sites for glutamylation, whereas some or all of the six residues (E435, E437, E439, E440, E441, and E442) in the β-tubulin CTT are likely the main sites for glycylation. This feature is the same as that of *Tetrahymena* (Redeker et al., 2005) but different from those of sea urchin sperm (Multigner et al., 1996), mammals (van Dijk et al., 2007), and *Drosophila* (Rogowski et al., 2009) in which the two species of tubulin likely undergo both glutamylation and glycylation.

We found that *TUA1(4A)* is almost completely immotile in contrast to *tpg1*, which displays motility albeit reduced. This may be due to the different degrees of tubulin polyglutamylation inhibition in their axonemes; *tpg1* retains polyglutamylated tubulin in the ciliary basal region presumably catalyzed by TTLL enzyme(s) other than TTLL9, while *TUA1(4A)* almost completely lacks polyglutamylation from the base to tip of the axoneme (Figure 2). Previously, we speculated that polyglutamylation increased inter-doublet friction of the outer-doublet microtubules. This speculation was based on the observation that the microtubule sliding in the axonemal fragments of *tpg1oda2* double mutant, which lacks outer-arm dynein, was faster than that in the axonemal fragments of *oda2*. We thought that the putative friction (strong dynein-microtubule interaction) might be important for the generation of ciliary motility (Kubo et al., 2010; 2012). The presence of motility in *tpg1* but not in *TUA1(4A)* cilia suggests that, if the inter-doublet friction was, in fact, important for motility, friction in the proximal region of the cilium may function as a trigger of the ciliary beating cycle. Another possible reason for the loss of motility in *TUA1(4A)* is that axonemal dyneins cannot generate force in the absence of the four glutamates in the α-tubulin CTT. Whether the loss of polyglutamylation per se is the cause of the motility loss in *TUA1(4A)* awaits further studies.

Motility analyses of *tpg1* combined with various dynein-lacking mutants led to a conclusion that polyglutamylation is important for the function of inner-arm dynein but not that of outer-arm dynein, because inner-arm dynein deficient mutants with normal outer-arm dynein, but not mutants lacking outer arm, remain motile in the absence of polyglutamylation (Kubo et al., 2010; 2012). A later detailed study strongly suggested that polyglutamylated tubulin directly associates with the nexin-dynein regulatory complex (N-DRC; Kubo et al., 2012; 2017), a structure that bridges between the adjacent outer-doublets and regulates the function of inner-arm dynein. However, our present study suggests that the polyglutamylated side chains or the glutamate residues of the α-tubulin CTT may also affect the function of outer-arm dynein since their loss resulted in a non-motile phenotype even when outer-arm dynein was intact. The true importance of the α-tubulin CTT needs to be examined using various dynein-lacking mutants in the background of mutations that inactivate polyglutamylation while preserving the four glutamates of α-tubulin CTT.

Many cells of *TUB2(Δ4E)* failed to swim (Figure 6A) though its axonemal structure was normal (Figure 8A). This suggests that negative charges of glutamate residues of the β-tubulin CTT are also involved in the ciliary motility. However, because the lack of glutamate residues in *TUB2(Δ4E)* coincidentally causes decreased glycylation (Figure 7), there is another possibility that glycylated tubulin also is affecting the ciliary motility. Indeed, tubulin polyglycylation is reported to regulate the function of axonemal dyneins (Gadadhar et al., 2021). Although “poly”glycylation likely does not occur in the *Chlamydomonas* axoneme, glycylation may be important for the ciliary motility. This can be clarified by generating a novel *Chlamydomonas* mutant lacking enzyme(s) responsible for glycylation.

In the *TUA1(4A)* axoneme, polyglutamylation was completely missing whereas glycylation increased (Figure 3C). In the *TUB2(Δ4E)* axoneme, in contrast, glycylation was missing whereas polyglutamylation slightly increased. This is consistent with the previous reports that there is an inverse correlation between the level of glutamylation and that of glycylation (Redeker et al., 2005; Wloga et al., 2009; Rogowski et al., 2009). In accordance with these studies, TTLL3 (monoglycylase) and TTLL7 (glutamylase) were found to compete for the same sites on β-tubulin *in vitro* (Garnham et al., 2017). This competition for the same substrate is interpreted as a regulatory mechanism to control the level of tubulin modification, but we still do not understand how the level of α-tubulin glutamylation affects the level of β-tubulin glycylation. Because polyglycylation was also found to modulate ciliary motility (Gadadhar et al., 2021), one idea is that polyglycylation can partially complement the motility defect caused by the lack of polyglutamylation. How these two modifications affect ciliary motility in a coordinated way still needs to be explored.

### Six glutamate residues of the β-tubulin CTT are dispensable for cell survival in Chlamydomonas

Unexpectedly, we found that the *TUB2(Δ6E)* cells are viable. This is different from *Tetrahymena* mutants with three mutations in five glycylation sites of β-tubulin (E437, E438, E439, E440, and E442) showing hypomorphic or lethal phenotype (Xia et al., 2000; Thazhath et al., 2002; Redeker et al., 2005). Although a *Chlamydomonas* cell appears to be inviable without the nine glutamates in the β-tubulin CTT (Figure 5B), our results indicate that the six glutamate residues (E435, E437, E439, E440, E441, and E442) corresponding to the five glycylation sites of *Tetrahymena* are dispensable for cell survival in *Chlamydomonas*.

In addition to this viable phenotype of *TUB2(Δ6E)*, we noticed some differences between the *TUB2(Δ6E)* cilia and those of the previously reported *Tetrahymena* mutants. The *Tetrahymena* mutants harboring a mutation in the glycylation sites of β-tubulin CTT partially lack the B-tubule outer doublets and the central-pair microtubules, and abnormally accumulate IFT-particle proteins between the ciliary membrane and outer doublets (Thazhath et al., 2002; Redeker et al., 2005). However, *TUB2(Δ6E)* cilium had normal outer doublets although it completely lacks the central apparatus (Figure 8A). In addition, IFT particles were of normal level or only slightly increased in the cilia (not shown), and bulged cilia reflecting the abnormal accumulation of IFT-particle proteins were not observed.

### Glutamate residues of the β-tubulin CTT may be involved in the IFT of tubulin

*TUB2(Δ6E)* is unable to regenerate cilia after pH-shock induced deciliation (Figure 6C). During the assembly of cilia, vast amounts of tubulins are transported into cilia by IFT (Craft et al., 2015). *In vitro* pull-down assay suggested that the N-terminal domains of IFT-particle proteins, IFT74 and IFT81, form a tubulin-binding module. More specifically, the calponin homology domain of the IFT81 N-terminus binds with the globular cores of the tubulin dimer and the basic IFT74 N-terminus likely associates with the acidic tail of β-tubulin (Bhogaraju et al., 2013). In accordance with this idea, the *Chlamydomonas* mutants deficient in either the IFT81 or the IFT74 N-terminus assemble flagella with reduced speed (Kubo et al., 2016; Brown et al., 2015). The inability of *TUB2(Δ6E)* to assemble full-length cilium is consistent with these previous results, indicating that the β-tubulin CTT is involved in the IFT of tubulin. To confirm this idea, the frequency of IFT in the *TUB2(D6E)* cilia should be investigated.

### Glutamate residues of the β-tubulin CTT may be involved in the katanin-mediated assembly of the central apparatus

The phenotype of *TUB2(Δ6E)* -having nonmotile cilia lacking the central apparatus -is shared by *pf15* and *pf19*, each with a mutation in a gene encoding a katanin subunit (Dymek et al., 2004; 2012). Katanin, an AAA ATPase (ATPases associated with diverse cellular activities), is one of the microtubule-severing enzymes (McNally and Vale, 1993). Recently, katanin was shown to be involved in the formation of the central apparatus by severing peripheral microtubules to produce central microtubule seeds (Liu et al., 2021). Furthermore, the β-tubulin CTT is necessary and sufficient for katanin to sever microtubules *in vitro* (Zehr et al., 2020). Therefore, the loss of the central apparatus in *TUB2(Δ6E)* lacking the β-tubulin CTT could also be due to the deficiency of katanin function. Even if this is the case, however, we still do not understand why, in the first place, such *Chlamydomonas* mutants with deficient katanin show normal cell proliferation because katanin has been suggested to be also involved in cell mitosis (Vale, 1991; McNally and Thomas, 1998; McNally et al., 2006). Therefore, various aspects of katanin function dependent on the β-tubulin CTT still need to be explored. Our system to introduce a tubulin mutation in *Chlamydomonas* must contribute to clarifying such important microtubule-based functions in the future.

## Materials and Methods

### Strains and Cultures

The cells (cc124 and generated mutants, Supplemental Table 1) were maintained on Tris-acetate-phosphate (TAP; Gorman and Levine, 1965) agar plates for long-term storage. For the experiments, the cells were cultured in liquid TAP medium at 25 °C with a light/dark cycle of 12:12 h and aeration.

### Generation of mutants by a CRISPR/Cas9 mediated gene editing

#### 1) Preparation of a Cas9/guide RNA ribonucleoprotein

The target sequences were searched using CRISPRdirect (http://crispr.dbcls.jp/). The crRNAs (Supplemental Table 2) and tracrRNAs (fasmac) were dissolved in Nuclease-Free Duplex Buffer (Integrated DNA technologies; IDT buffer), dispensed in micro-8-tube strips, and cryo-preserved at -80 °C. The crRNA (1 µl, 40 µM) was annealed with tracrRNA (1 µl, 40 µM) by heating at 95 °C for 2 min and then cooling down gradually to 25 °C over 90 min. 2 µl of annealed guide RNA was then incubated with 0.5 µl Cas9 protein (10 µg/ml; fasmac) with 7.5 µl IDT buffer at 37 °C for 15 min to generate Cas9/gRNA RNP.

#### 2) Transformation of the cell

The cells were grown on a TAP agar plate for 5 days under constant light and treated with ∼6 ml autolysin (Picariello et al., 2020) for an hour to remove cell walls. The cells were then agitated gently at 40 °C for 30 min. The cells were collected by centrifugation (∼2,000 rpm, 3 min, room temperature) and washed by 2% sucrose in the TAP medium (TAP sucrose). After centrifugation (∼2,000 rpm, 3 min, room temperature), the cells were resuspended to a concentration of 2-7×10^9^ cells/ml. Cas9/gRNA RNP (10 µl), donor DNA (∼10 µl; up to 2 µg; Table S2), and the cells (∼110 µl) were mixed to give a final volume of 125 µl. The mixture was transferred to a cuvette (0.2 cm gap; Bio-Rad) and electroporated (ECM 630 Electroporation System, BTX) immediately at 350 V, 25Ω, and 600 µF. The cuvette was then incubated at 15 °C for 60 min. The transformed cells were suspended into 10 ml of TAP sucrose and gently rocked for 24 hours under dim light. The cells were collected by centrifugation, plated on a TAP-agar containing 10 µg/ml hygromycin or 10 µg/ml paromomycin, and cultured for 5-7 days under constant light.

### Western blotting

Western blotting was performed according to Towbin et al. (1979) with some modifications. SDS-protein samples were separated by electrophoresis using a handmade SDS-polyacrylamide gel (7.5 or 9% polyacrylamide) and transferred to a PVDF membrane (Immobilon-P, pore size 0.45 µm; Merck-Millipore). The membrane was probed with primary antibodies listed in Supplemental Table 3.

### Production of the antibodies against the central-apparatus proteins

The cDNA sequences encoding partial CPC1 (Cre03.g183200; amino acids 395-801), FAP221 (Cre11.g476376; amino acids 1-762), FAP101 (Cre02.g112100; amino acids 652-835), KLP1 (Cre02.g073750; amino acids 451-633), PF20 (Cre04.g227900; amino acids 9-606), and FAP7 (Cre12.g531800; amino acids 269-507) were inserted into the pGEX vector and the polypeptides were expressed in BL21 (DE3) competent E. coli (NEB). Rabbit polyclonal antibodies were raised against the purified polypeptides.

### Isolation of cilia

Cilia were isolated according to Witman et al. (1972) with some modifications. Briefly, fully grown cells were collected with centrifugation (3,000 rpm, 5 min) and treated with 1 mM dibucaine-HCl (Wako) to amputate their cilia. Cilia were washed and collected by centrifugation (10,000 rpm, 20 min, 4 °C). Occasionally, flagella were demembranated by 0.1% Igepal CA-630 (Sigma) to prepare axonemal samples.

### Indirect immunofluorescence microscopy

Immunostaining was carried out following Sanders and Salisbury (1995). Cells from 5-10 ml culture were collected by gentle centrifugation and treated with 6 ml autolysin for 60 min to remove cell walls. The cells were washed with NB buffer (6.7 mM Tris-HCl (pH 7.2), 3.7 mM EGTA, 10 mM MgCl2, and 0.25 mM KCl) and placed on an 8 well-slide glass (8 mm well; Matsunami) treated with polyethylenimine. The cells were then fixed with -20 °C methanol and -20 °C acetone for 5 min, respectively. The cells were first treated with blocking buffer (1% BSA and 3% Fish Skin gelatin in PBS) and incubated with primary antibodies (Supplemental Table 3) followed by secondary antibodies (goat anti-rabbit IgG Alexa 488, 1:200, Invitrogen; goat anti-mouse IgG Alexa Fluor 594, 1:200, Invitrogen). The cells were treated with antifade mountant (SlowFade Diamond, Thermo Fisher Scientific) and encapsulated with a glass cover slip. The sample was examined with a microscope (BX53, Olympus) equipped with an objective lens (100× UPlan FL N Oil objective, Olympus). Images were acquired with a CCD camera (ORCA-Flash4.0 sCMOS, Hamamatsu).

### Transmission electron microscopy (TEM)

TEM observation of the isolated axonemes was performed as described previously (Huang et al., 1979). Briefly, the isolated axonemes were fixed for 1 hour with 2% glutaraldehyde in 10 mM phosphate buffer (pH 7.0) containing 1% tannic acid. After three washes with the buffer, the axonemal pellets were fixed on ice for 45 min with 1% osmium tetroxide in phosphate buffer. The pellets were dehydrated with graded series of ethanol. The pellets were treated with propylene oxide and then embedded in epoxy resin. Thin sections were made by ultra-microtome and were stained with uranyl acetate and lead citrate. Images were obtained by JEM-2100F transmission electron microscope (JEOL, Tokyo, Japan).

### Ciliary length measurement and motility analyses

To assess ciliary regeneration kinetics, cells were deflagellated by pH shock (Rosenbaum et al., 1969). After the deflagellation, aliquots of the cells were isolated and fixed by 1% glutaraldehyde at 15 min intervals up to 180 min. One cilium each from at least 30 cells was measured by ImageJ to obtain the average length of each time point.

Swimming velocity was acquired by tracking images of the moving cells. Briefly, the cells under the dark-field microscope equipped with 40x objective were recorded using a digital camera, and the obtained movies were processed with ImageJ.

Microtubule sliding velocity during axonemal disintegration was measured as described previously (Kurimoto and Kamiya, 1991). Briefly, fragmented axonemes (∼5 µm in length) were placed in a perfusion chamber under a dark-field microscope. Microtubule sliding was induced by a solution containing 0.5 µg/ml of Type VII bacterial protease (Sigma) and ATP. This process was recorded using a ×100 objective, an oil-immersion dark-field condenser, a light source of a mercury lamp (Olympus, U-RLF-T), and a CCD camera (Olympus, M-3204C).

## Acknowledgment

We greatly appreciate Dr. Ritsu Kamiya (Chuo University, Tokyo) for critically reading the manuscript and a constructive discussion.

## Competing interests

No competing interests declared.

## Funding

This work was supported by Takeda Science Foundation (to T.K. and T.O.), the Uehara Memorial Foundation (to T.K.), Koyanagi Foundation (to T.K.) and Japan Society for the Promotion of Science [19K16123 (to T.K.), 21H02654 (to T.O.), and 21H05248 (to M.K.)]

---

**Supplemental Figure 1.**
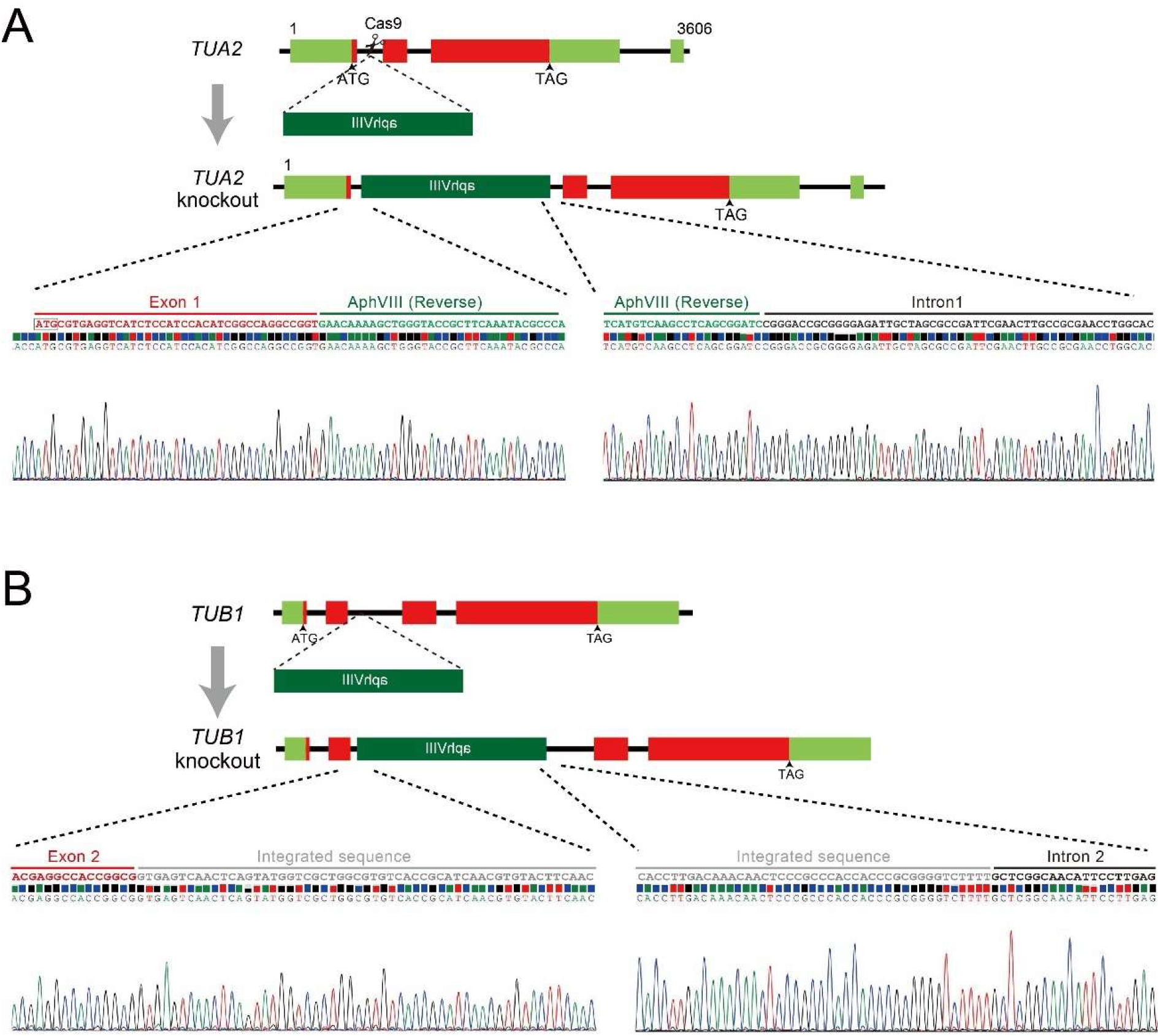
Generation of *tua2(int1)* and *tub1(int2)* mutants. Production of (A) *tua2 (int1)* and (B) *tub1(int2)* by a CRISPR/Cas9 mediated gene editing. A paromomycin-resistant gene cassette (aphVIII) was inserted into either intron 1 of *TUA2* or intron 2 of *TUB1*. Sequencing results of the integrated areas are shown.

**Supplemental Figure 2.**
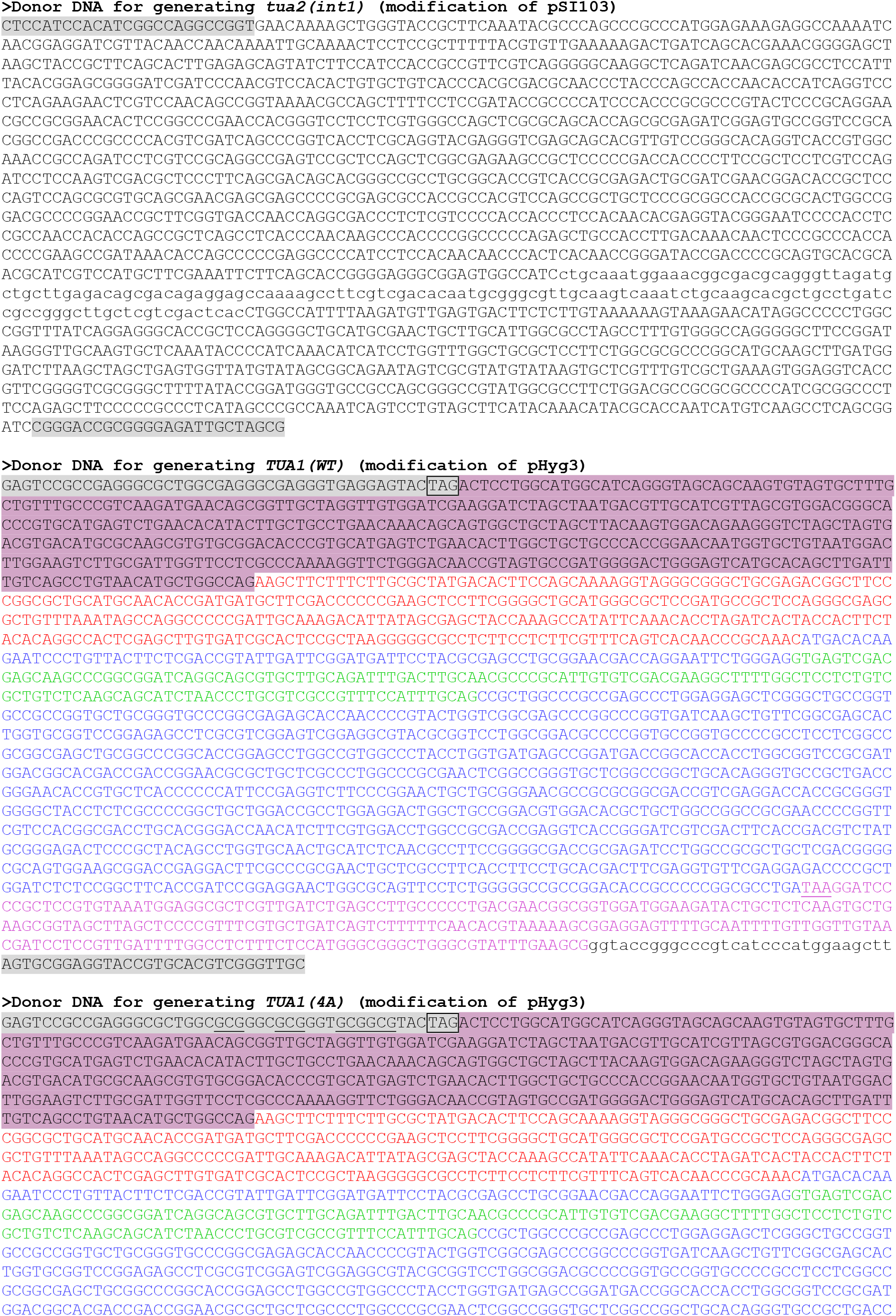
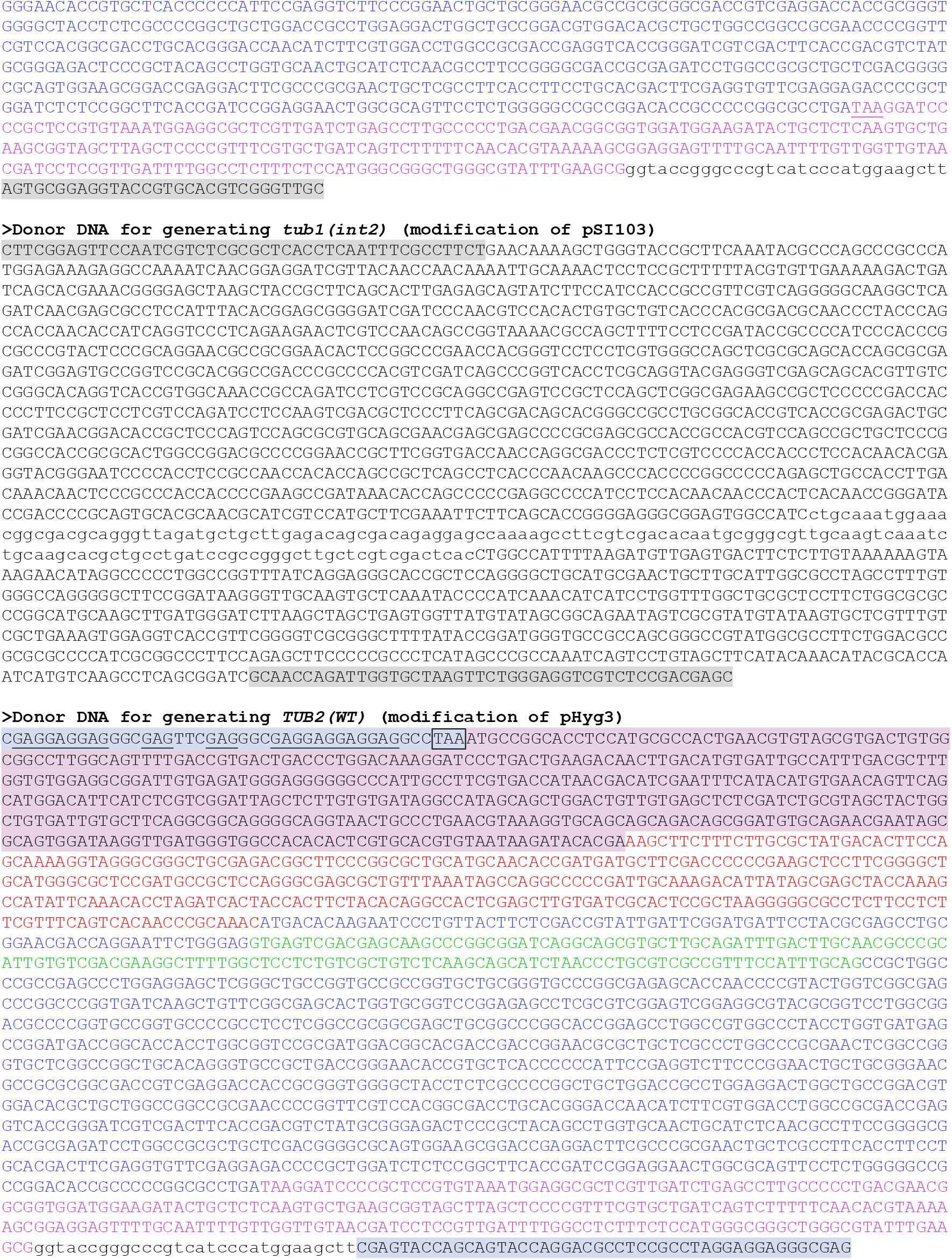
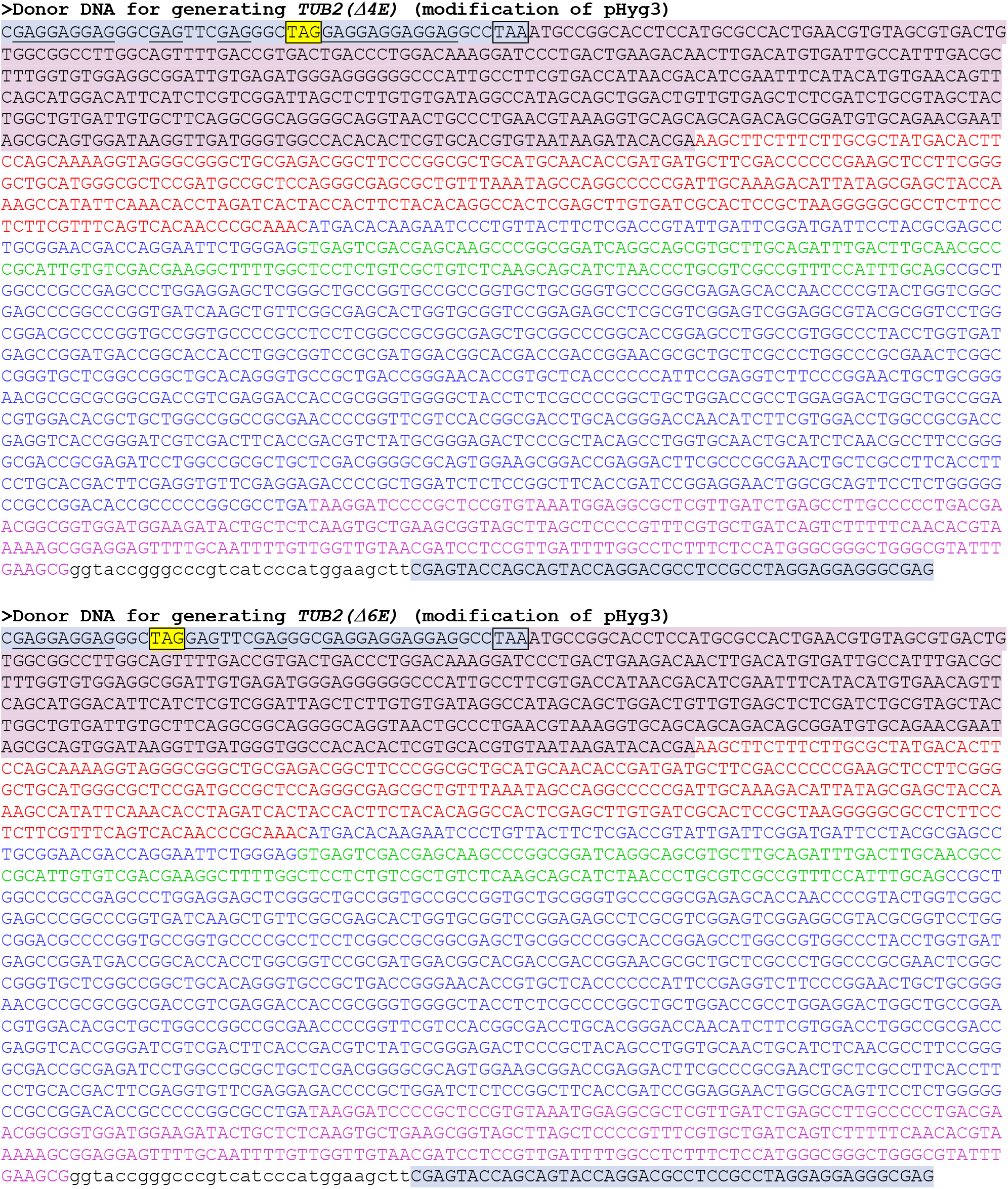

**Supplemental Figure 3.**
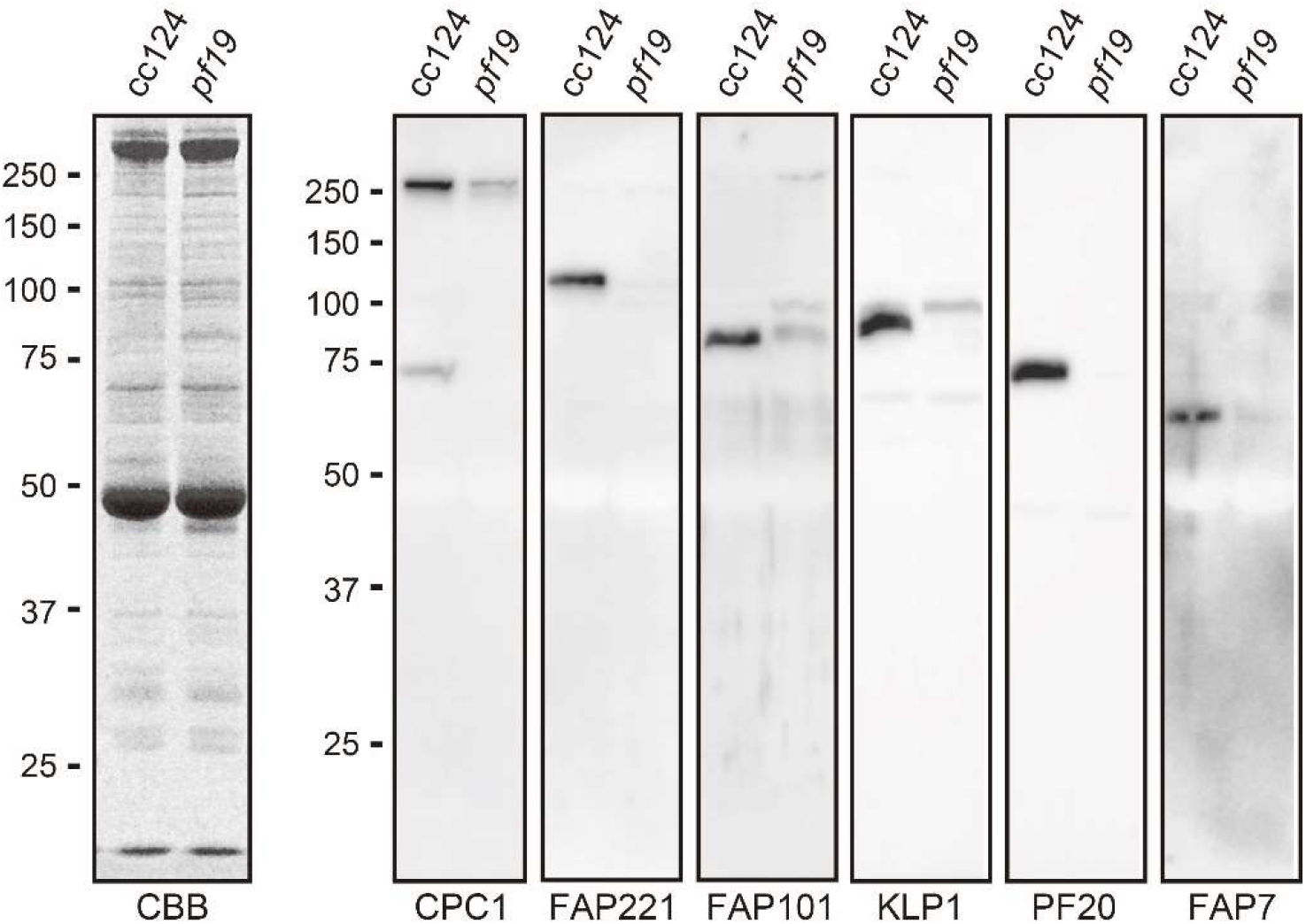
Characterization of the polyclonal antibodies against the central-apparatus proteins. CBB-stained gel (left) and Western blotting (right) of the axonemes of wild-type (cc124) and *pf19* deficient in the central apparatus. Purified antibodies against CPC1 (205.1 kDa), FAP221 (104 kDa), FAP101 (85.7 kDa), KLP1 (83 kDa), PF20 (53 kDa), and FAP7 (54.7 kDa) were examined.

**Supplemental Table 1.** List of mutants generated in this study

| **Name** | **Origin** | **Mating type** | **Antibiotic resistance** |
| --- | --- | --- | --- |
| *tua2(int1)* | cc124 | Minus | Paromomycin |
| *TUA1(WT)* | *tua2(int1)* | Minus | Hygromycin, paromomycin |
| *TUA1(4A)* | *tua2(int1)* | Minus | Hygromycin, paromomycin |
| *tub1(int2)* | cc124 | Minus | Paromomycin |
| *TUB2(WT)* | *tub1(int2)* | Minus | Hygromycin, paromomycin |
| *TUB2(Δ4E)* | *tub1(int2)* | Minus | Hygromycin, paromomycin |
| *TUB2(Δ6E)* | *tub1(int2)* | Minus | Hygromycin, paromomycin |

**Supplemental Table 2.**
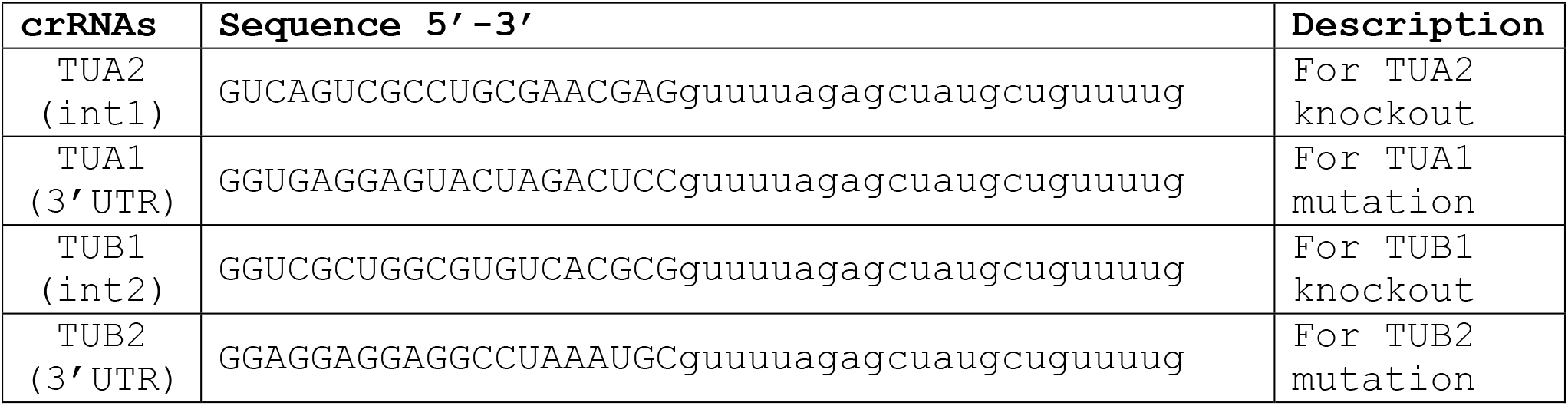
List of crRNA sequences

**Supplemental Table S3.**

| Antibody (clone #) | Host | Dilution | Reference |
| --- | --- | --- | --- |
| Anti-α-tubulin (B512) | Mouse | 1:5000 (WB) | Sigma-Aldrich |
| Anti-polyglutamylated tubulin (polyE#1) | Rabbit | 1:2000 (WB) | Kubo et al. (2017) |
| Anti-polyglutamylated tubulin (polyE#2) | Rabbit | 1:2000 (WB),<br>1:200 (IFM) | Kubo et al. (2017) |
| Anti-glutamylated tubulin (GT335) | Mouse | 1:2000 (WB),<br>1:200 (IFM) | Adipogen Life Sciences |
| Anti-glycylated tubulin (Gly-pep1) | Rabbit | 1:2000 (WB),<br>1:200 (IFM) | Funakoshi |
| Anti-CPC1 | Rabbit | 1:5000 (WB) | This study |
| Anti-FAP221 | Rabbit | 1:5000 (WB) | This study |
| Anti-FAP101 | Rabbit | 1:5000 (WB) | This study |
| Anti-KLP1 | Rabbit | 1:5000 (WB) | This study |
| Anti-PF20 | Rabbit | 1:5000 (WB) | This study |
| Anti-FAP7 | Rabbit | 1:5000 (WB) | This study |

## Notes

### Competing Interest Statement

The authors have declared no competing interest.

